# A Platform for Mitochondrial Profiling in Enriched Kidney Segments Under Thermodynamic Control in Mice and Humans

**DOI:** 10.1101/2025.05.05.652276

**Authors:** Stephen T Decker, Precious C Opurum, Ran Hee Choi, Venisia L Paula, Za’rya T Smith, Anu S Kurian, Deborah Stuart, Linda Nikolova, Alejandro Sanchez, Suguru Takayama, Laith Al-Rabadi, Nirupama Ramkumar, Kelsey H Fisher-Wellman, Katsuhiko Funai

**Affiliations:** Center for Metabolic Health, University of Utah, Salt Lake City, Utah; Department of Nutrition and Integrative Physiology, University of Utah, Salt Lake City, Utah; Department of Kinesiology, University of Utah, Salt Lake City, Utah; Haumana’O Pasifika Program, University of Utah, Salt Lake City, Utah; Division of Nephrology and Hypertension, University of Utah School of Medicine, Salt Lake City, Utah; Electron Microscopy Core Facility; University of Utah; Salt Lake City, UT; USA; Division of Urology, Department of Surgery, University of Utah School of Medicine, Salt Lake City, Utah, USA; Huntsman Cancer Institute, Cancer Hospital, Salt Lake City, Utah, USA; Department of Cancer Biology, Wake Forest University School of Medicine, Winston-Salem, North Carolina, USA

## Abstract

Mitochondrial function varies widely across kidney nephron segments, yet conventional approaches lack the resolution and control needed to assess cell-type-specific bioenergetics in situ. We present a methodological platform that enables segment-resolved profiling of mitochondrial respiration, conductance, and membrane potential in freshly isolated mouse and human nephron segments. Combining mechanical sieving and adhesion-based enrichment with permeabilized high-resolution respirometry, we adapted the creatine kinase clamp to quantify oxygen flux and mitochondrial membrane potential across defined free energies. Using this approach, we found that proximal tubules exhibit high respiratory conductance and dynamic mitochondrial polarization, while distal tubules and glomeruli maintain static membrane potential and low conductance. In a model of adenine-induced nephropathy, only proximal tubule mitochondria showed marked reductions in respiration and ATP production. This segment-specific dysfunction was not detectable in bulk mitochondrial isolates. We then demonstrate that these methods can be applied in human kidney samples. Our approach provides thermodynamically anchored, segment-resolved insight into mitochondrial adaptation under physiological and pathological conditions. It is broadly applicable to other tissues with metabolic heterogeneity and compatible with disease models, genetic tools, and pharmacological interventions. This platform bridges a critical gap between conventional respirometry and functional mitochondrial phenotyping in native tissue structures.

## Introduction

Kidneys are the second-most bioenergetically active organ per unit mass, surpassed only by the heart^1–7^. This high metabolic activity is required to support the nephron’s critical roles in filtration, reabsorption, and excretion. Each day, the kidneys process approximately 130 liters of glomerular filtrate, reabsorbing nearly 99% of this volume back into circulation and excreting only 1–2 liters as urine^8,9^. This extraordinary resorptive capacity is driven by ATP-dependent transport systems that operate continuously to move ions and solutes against their electrochemical gradients^8,10,11^. Key drivers of this active transport include Na^+^/K^+^-ATPase^10,12^, Ca^2+^/Mg2+-ATPase^13^, and vacuolar H^+^-ATPase^14,15^, which consume significant amounts of ATP to maintain electrolyte and fluid homeostasis.

Mitochondria are central to meeting these energy demands. As the primary generators of ATP via oxidative phosphorylation, mitochondria are highly dynamic organelles capable of adapting structurally and functionally to the cell’s metabolic state^16,17^. In response to energetic demands, mitochondria undergo changes in cristae architecture^18,19^, fission and fusion dynamics, and the expression and activity of electron transport chain (ETC) components^20–22^. These adaptations enable fine-tuning of mitochondrial efficiency, capacity, and coupling, supporting tissue-specific metabolic requirements.

Within the nephron, mitochondrial morphology, density, and function vary substantially by segment, reflecting the physiological heterogeneity of renal epithelial cells^7^. Proximal tubules (PT), which are responsible for reabsorbing approximately 65% of the glomerular filtrate^10^, exhibit the highest mitochondrial density and possess elaborate cristae networks to support their intense ATP requirements^1,23,24^. In contrast, distal tubules (DT) and glomerular cells (Glom), which are involved in fine-tuning solute handling and maintaining the filtration barrier, respectively, exhibit lower mitochondrial content and simpler ultrastructure^23,24^. These segmental differences have important implications for susceptibility to metabolic stress and injury, with the PT being particularly vulnerable due to its high baseline metabolic rate and limited energy reserve^23–25^.

Despite the recognized heterogeneity in nephron bioenergetics, most existing approaches to studying mitochondrial function in the kidney rely on mitochondria isolated from whole-kidney homogenates or cultured renal cell lines^1,26,27^. These models lack the resolution to detect segment-specific functional differences and are often poorly suited to capturing mitochondrial responses under physiologically relevant conditions. Whole-kidney mitochondrial isolations, while applicable for bulk assessments^28^, blend populations from diverse nephron segments and may selectively bias the mitochondrial population^29,30^, thereby diluting or masking localized dysfunction. Meanwhile, cultured cell models often fail to recapitulate in vivo cell polarity, tissue architecture, and the integrated stress responses that define renal physiology^26,27,31^. For example, the proximal tubule is known to be a primary target in toxicological models such as the high-adenine diet (AD), which induces fibrosis and tubular injury^32–34^; however, these effects are often obscured in whole-kidney preparations or incompletely modeled in vitro^35^.

To overcome these limitations, we developed a comprehensive methodological platform to enrich and interrogate mitochondria from specific nephron segments using a combination of mechanical sieving and adhesion-based separation^36,37^. Here, we apply this approach to evaluate how PT, DT, and Glom mitochondria respond to basal conditions and to nephrotoxic stress induced by an AD in mice, and confirm that these methods can be applied to investigate segment-enriched mitochondrial function in humans. We demonstrate that proximal tubules exhibit high bioenergetic adaptability while distal nephron segments and glomeruli maintain static, tightly polarized mitochondria. This platform enables segment-resolved mitochondrial profiling in complex tissues and offers a generalizable strategy for investigating sub-organ energetics in physiology and disease.

## Results

### Segment-Resolved Isolation Enables Mitochondrial Analysis in Native Nephron Structures

Standard methods for assessing mitochondrial function generally rely on mitochondria isolated from whole kidney tissue (often cortex-only preparations) or cultured renal cell lines. While these approaches are widely used and beneficial, they lack the sensitivity required to detect subtle, segment-specific mitochondrial changes resulting from in vivo interventions. To address these limitations, we employed a sieving and adhesion-based enrichment method (Figure 1A) to isolate distinct nephron segments – PT, DT, and Glom – from freshly harvested mouse kidneys^36,37^. We then ran a standard SUIT protocol on these samples to determine site-specific mitochondrial respiration at (Figure 1B).

**Figure 1.**
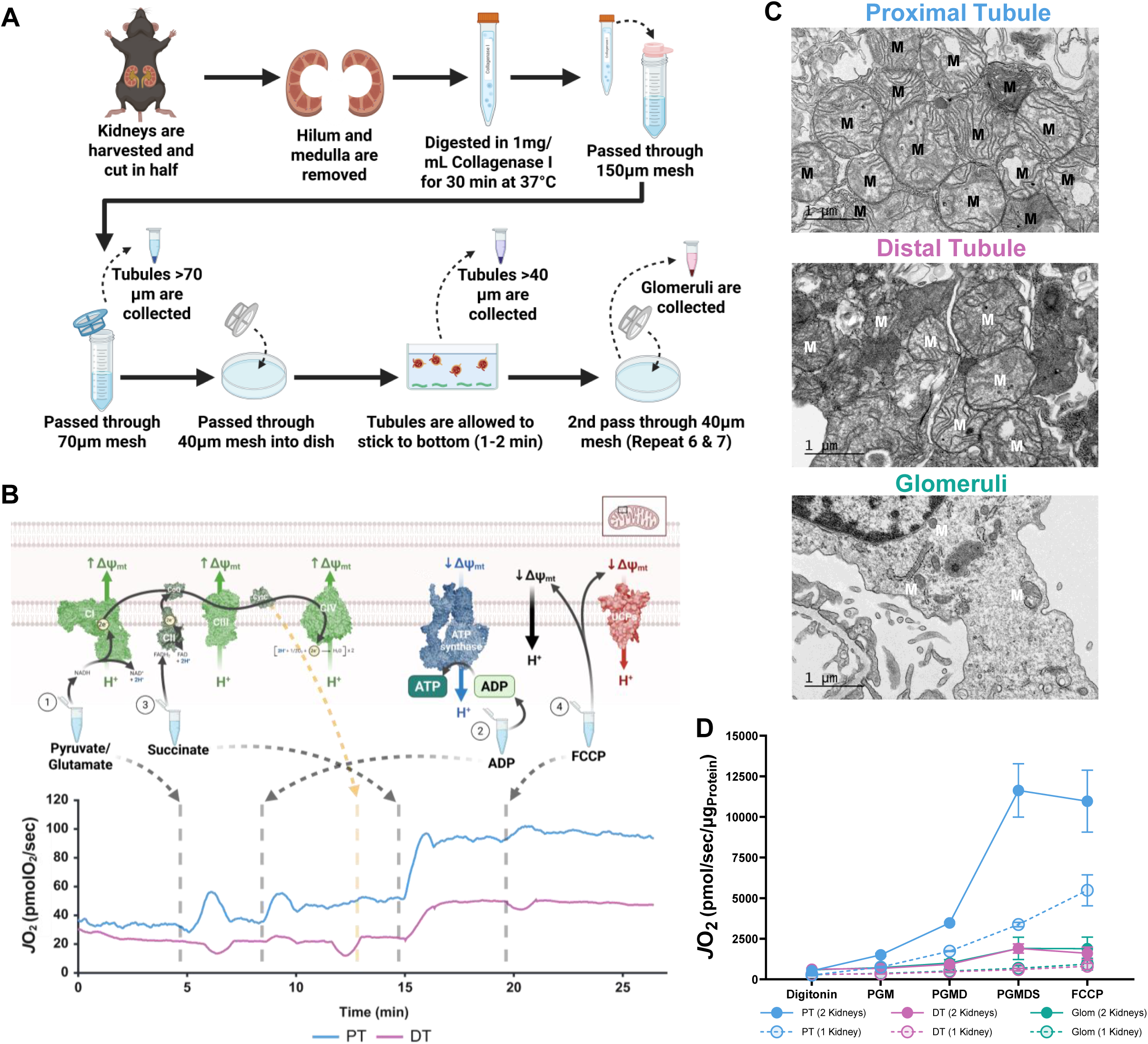
A) Schematic outlining the sieving procedure in kidneys harvested from mice. B) Protocol outline of the substrate-uncoupler-inhibitor titration (SUIT) protocol. Substrates, which activate the electron transport chain, are added sequentially. Pyruvate and glutamate produce NADH^+^ via the TCA cycle, which is used as a substrate for Complex I (CI), and in the absence of ADP, represents a “leak” state. ADP is then added to the chambers to activate the production of ATP, thus representing a phosphorylation state. Cytochrome C (CytC), an indicator of membrane integrity, and succinate, a substrate for complex II (CII), are then added to assess CI-and CII-dependent respiration. FCCP is added to assess supramaximal ETC activity. C) Electron microscopy images of the sieved tissues, indicating the abundance and morphological differences between proximal tubule, distal tubule, and glomerular mitochondria (marked by “M”). D) Initial pilot experiments showing that mitochondrial respiration can be measured in enriched samples from mice using both harvested kidneys or a single harvested kidney. *P*: Pyruvate; *G*: Glutamate; *M*: Malate; *D*: ADP; *S*: Succinate; *FCCP*: Carbonyl cyanide 4-(trifluoromethoxy)phenylhydrazone

In brief, kidneys were harvested, the medulla was carefully removed, and the cortical tissue was finely diced to facilitate enzymatic digestion. Tissue digestion was carried out using collagenase type I, followed by sequential filtration through cell strainers with decreasing mesh sizes (100 μm, 70 μm, and 40 μm). PT segments were selectively retained on the 70 μm mesh, while DT and Glom segments passed through and were subsequently collected on the 40 μm mesh. We leveraged their differential adhesive properties to separate DT from Glom further; DT segments adhered to culture dishes, whereas Glom remained suspended. The isolated DT segments were gently scraped from the dishes, and the suspended Glom were separately collected. Transmission electron microscopy confirmed the preservation of mitochondrial ultrastructure within each fraction, with PT samples displaying densely packed mitochondria with well-defined cristae, while DT and Glom exhibited sparser and morphologically simpler mitochondria (Figure 1C). Segmental enrichment was validated by quantitative PCR for nephron-specific markers: Nphs1 (Glom), Slc5a2 (PT), and Aqp2 (DT) (Figure S1).

To evaluate mitochondrial function in these enriched nephron segments, we utilized a standardized substrate-uncoupler-inhibitor titration (SUIT) protocol in permeabilized preparations^38^. This approach allowed comprehensive measurement of mitochondrial respiration using HRR (Oroboros O2K) in conjunction with fluorometric assessments (Fluorolog-QM)^39,40^. SUIT protocols systematically evaluate ETC functionality by quantifying oxygen consumption rates (*J*O_2_) as specific substrates are sequentially oxidized through Complexes I to IV (Figure 1B)^38,41^. This system has the advantage over other systems that have been previously utilized^42,43^, as this system allows for the measurement of *J*O_2_ in nonadherent, intact systems, allows for the rapid diffusion of O_2_ via continuous stirring (thereby increasing the diffusion of O_2_ in the media and increasing the sensitivity of the O_2_ flux measurements), customizable SUIT protocols, and continuous, real-time measurement of O ^41^.

Notably, we demonstrated that reliable mitochondrial respiration measurements could be achieved with as little as approximately 9 μg of protein per respirometry chamber, as illustrated in initial pilot experiments combining both kidneys from a single mouse (Figure 1D). We conducted additional studies using a single kidney to assess whether this method could be applied to more limited sample volumes, such as when only one kidney is available due to concurrent tissue requirements for other analyses (e.g., histology, PCR, or mass spectrometry). These experiments confirmed that the assay could be successfully performed using one kidney, albeit with some increased technical variability due to lower tissue yield (Figure 1D).

### Adenine Diet Selectively Impairs Mitochondrial Function in Proximal Tubules

To test the sensitivity of our platform to physiologically relevant injury, we exposed mice to an AD, a well-established model of tubulointerstitial nephropathy that predominantly targets the proximal tubule^32–34,44,45^. Mice were fed alternating concentrations of 0.15% and 0.25% adenine over a two-week cycle to balance toxicity with survival (Figure 2A). AD-fed mice exhibited significant reductions in body weight, kidney mass, and liver mass compared to controls (Figure 2B, S2A-B), consistent with systemic catabolic effects. Histological analysis confirmed extensive cortical fibrosis, including tubular atrophy and interstitial expansion (Figure 3C). Quantitative image analysis using an in-house pipeline developed in Python revealed increases in tubulointerstitial fibrosis and increases in tubular luminal area, indicating widespread tubular damage (Figure S3). However, in agreement with previous literature, there were only modest changes in glomerular corpuscle area, fibrotic area, Bowman’s space area, and tuft area (Figure S3), indicating that the adenine diet robustly impacts tubules, but not glomeruli^32–34,44^.

**Figure 2.**
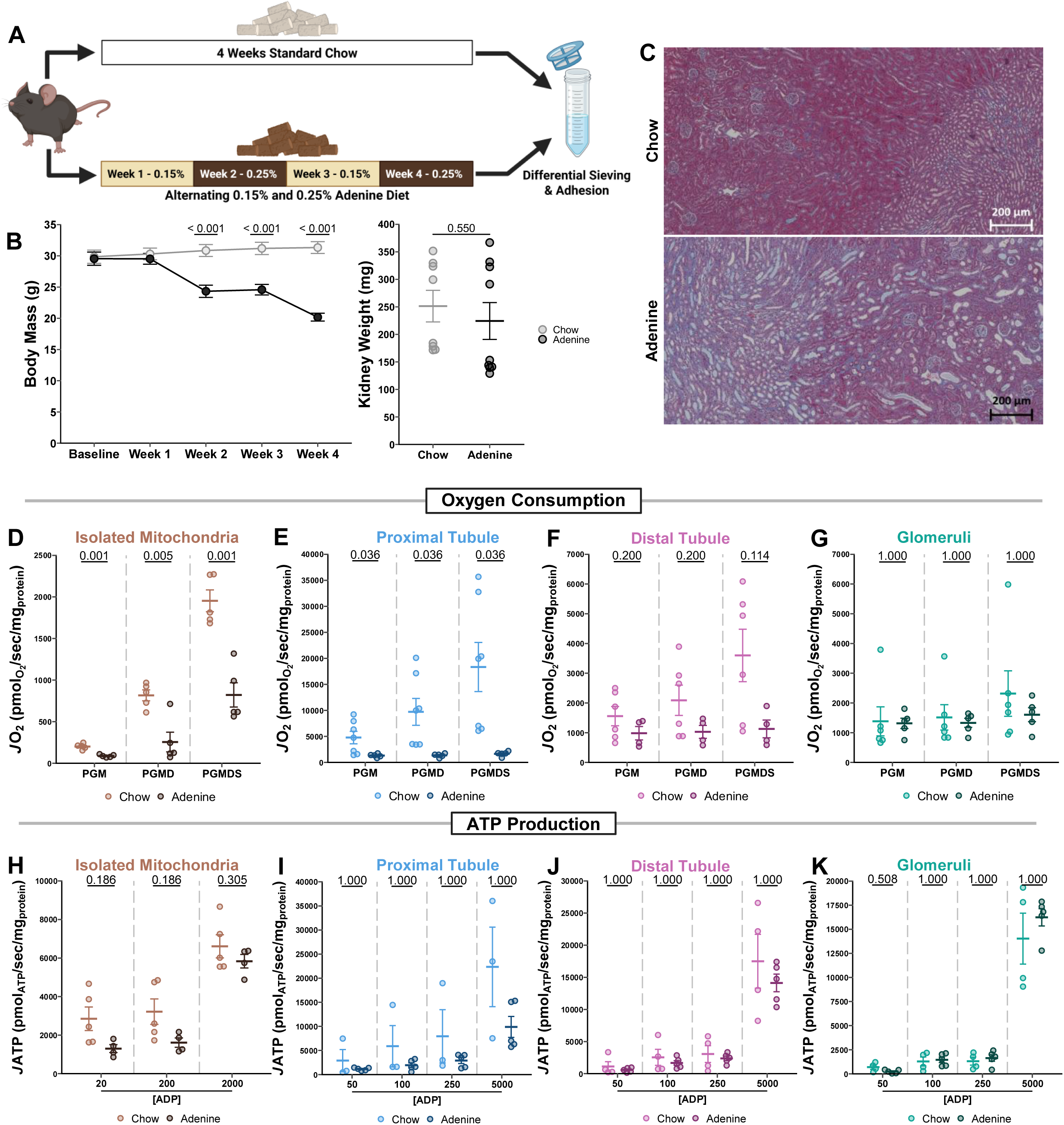
A) A schematic outlining the adenine diet protocol. B) Body weights over the course of the study, and kidney weights at the time of euthanasia. C) Repeasentative Masson’s Trichrome images of the kidneys of chow-fed and adenine-fed mice. D-G) Oxygen consumption rates (*J*O_2_) of mitochondria isolated from whole kidney (D), and enriched proximal tubule (E), distal tubule (F), and glomerular samples (G). H-K) ATP production rates (*J*ATP) of mitochondria isolated from whole kidney (H), and enriched proximal tubule (I), distal tubule (J), and glomerular samples (K). *P*: Pyruvate; *G*: Glutamate; *M*: Malate; *D*: ADP; *S*: Succinate.

**Figure 3.**
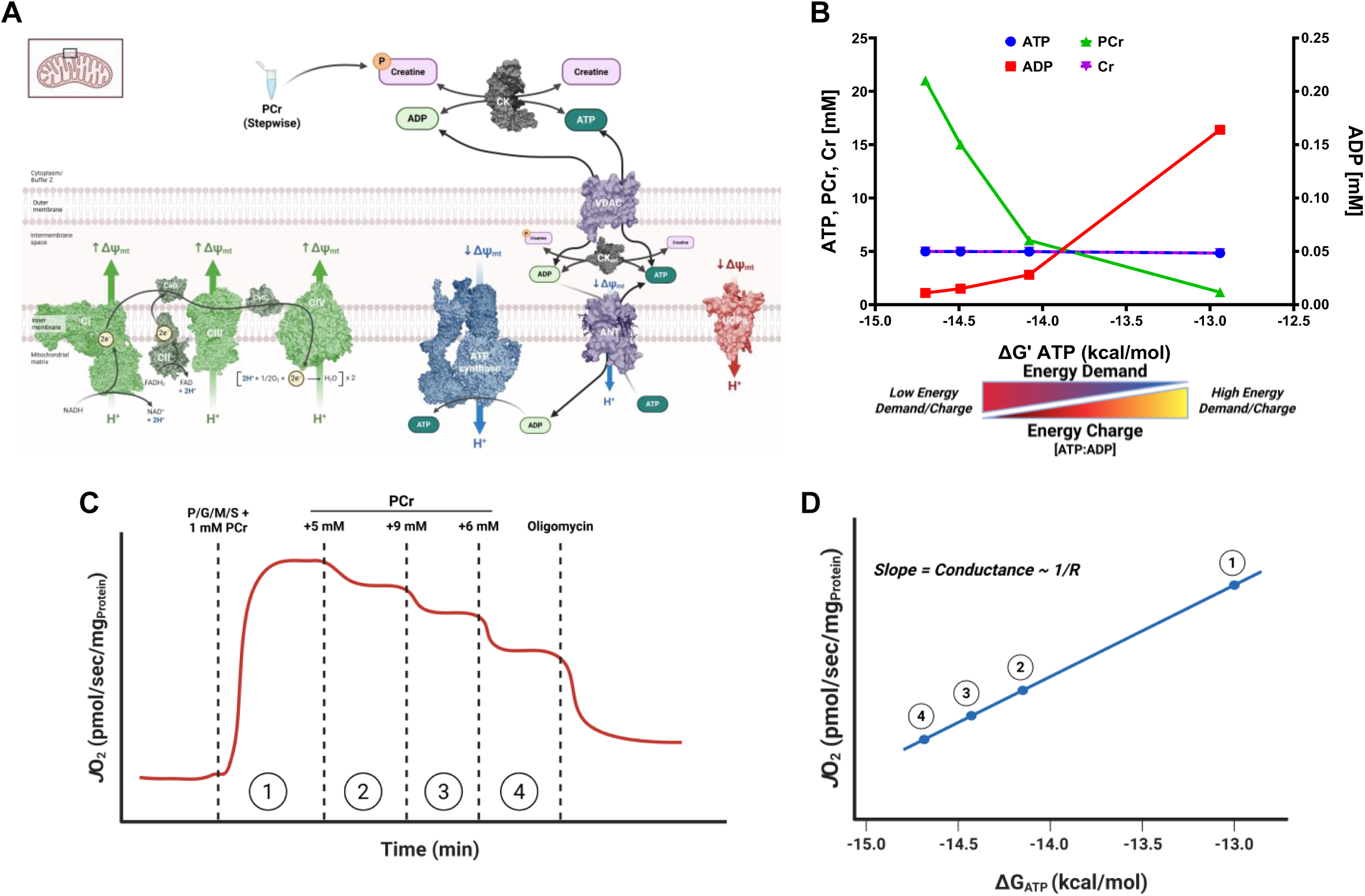
Shematics outlining the foundational concept of the creatine kinase (CK) clamp assay. The CK clamp assay works via modulating the ADP/ATP ratio under controlled bioenergetic conditions using standard substrates and increasing phosphocreatine (PCr) concentration (A). The use of PCr in the CK clamp allows for the controlled production of ATP under stable ADP concentations, and further allows for the calculation of the Gibbs free energy of ATP synthesis (ΔG’_ATP_), providing more detailed interrogation of bioenergetic properties of mitochondria (B). The addition of PCr under these standard conditions will act as a bioenergetic “brake” on the mitochondria, slowing the respiration rate as the [PCr] increases (C). Therefore, the controlled bioenergetic state of the mitochondria can be used to estimate the conductance of the electron transport chain via the slope of the relationship between oxygen consumption (*J*O_2_) and ΔG’_ATP_ (D).

To establish a functional benchmark, we first isolated mitochondria from whole kidney cortex of chow-and AD-fed mice using conventional differential centrifugation. Consistent with prior reports, AD feeding led to a robust ∼75% reduction in *J*O_2_ in isolated mitochondrial preparations (Figure 2D), confirming global mitochondrial impairment. However, this bulk measurement masks cellular heterogeneity. We applied our sieving/adhesion method and performed high-resolution respirometry on permeabilized PT, DT, and Glom segments. We observed a striking ∼90% reduction in state III *J*O_2_ in PT, a ∼75% reduction in DT, and a non-significant decrease in Glom (Figure 2D-G). These data confirm the segment-specific vulnerability of PT and DT to AD-induced mitochondrial dysfunction and reinforce the relative resistance of glomerular cells to AD-induced injury.

To assess the downstream bioenergetic consequences of these respiratory changes, we quantified ATP production using an NADPH-based fluorescence assay^40,46,47^. Interestingly, while isolated mitochondria and DT segments from AD-fed mice retained the ability to generate ATP, PT exhibited a significant and disproportionate reduction in ATP production (Figure 2H-K). This suggests that DT mitochondria may retain sufficient inner membrane potential and proton motive force to sustain ATP synthesis despite compromised oxygen consumption, consistent with other observations^23,24^. In contrast, PT mitochondria are more severely compromised, likely reflecting damage to multiple components of oxidative phosphorylation machinery, leading to both impaired respiration and ATP generation^44,45^.

### CK Clamp Reveals Segment-Specific Differences in Respiratory Conductance

While traditional SUIT protocols quantify mitochondrial respiration, they lack the thermodynamic context to assess efficiency or adaptability under dynamic energetic loads. Specifically, SUIT protocols do not allow for quantifying key bioenergetic variables such as the Gibbs free energy of ATP synthesis (ΔG_ATP_) or the conductance of electrons through the ETC. As a result, they offer limited mechanistic resolution regarding the efficiency and control of oxidative phosphorylation.

To overcome these limitations, we implemented the creatine kinase (CK) clamp^3,47–50^, a technique that enables fine control of cellular energy charge through the use of phosphocreatine (PCr) and exogenous CK enzyme. This method emulates dynamic changes in energy demand by systematically titrating PCr while maintaining constant levels of ATP and ADP through the action of CK. This system closely mimics physiological transitions between energetic states— ranging from rest (high PCr, low ADP) to high demand (low PCr, high ADP)—and allows for the calculation of respiratory sensitivity to ATP free energy across this gradient (Figure 3A)^47,49,50^. The CK clamp relies on the reversible reaction catalyzed by CK: PCr + ADP ⇌ Cr + ATP. This equilibrium buffers changes in ADP concentration and creates stable energetic states. As PCr increases, ATP is regenerated more readily, reducing the need for mitochondrial ATP synthesis and thereby decreasing *J*O2 (Figure 3B-C). Because the electrochemical proton gradient drives mitochondrial ATP production, the titration of PCr modulates ΔG_ATP_ in a controlled manner, which reflects the driving force for oxidative phosphorylation. By plotting *J*O_2_ against ΔG_ATP_, we generate a linear force-flow relationship, the slope of which represents the conductance of the ETC (Figure 3D)^47,48^.

Using this approach, we first assessed the response of mitochondria isolated from whole kidney cortex. These preparations demonstrated robust modulation of *J*O_2_ in response to PCr titration, and the AD induced a marked reduction in both oxygen consumption and conductance, confirming a loss of mitochondrial bioenergetic efficiency (Figure 4A-B). These results aligned well with those from SUIT-based assays and served as a baseline for comparison to enriched nephron segment samples. We next applied the CK clamp to enriched PT, DT, and Glom segments obtained using the sieving and adhesion technique. Among these, only PT showed a consistent and significant reduction in respiration in response to the AD (Figure 4C-H). This observation reinforces the conclusion that PTs are the primary targets of bioenergetic dysfunction in this model of renal injury, while DT and Glom remain comparatively resistant.

**Figure 4.**
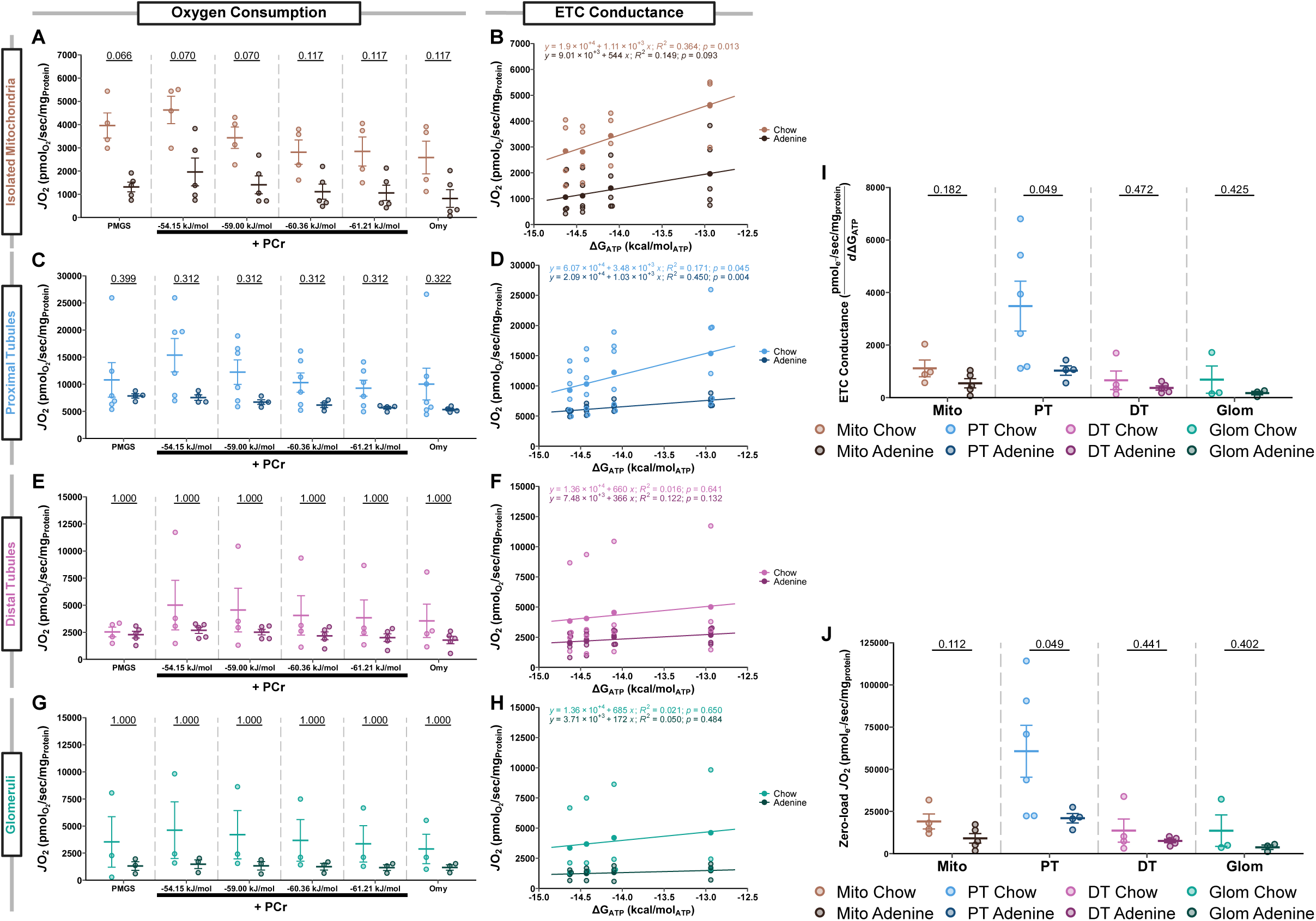
Results of the CK clamp assay on oxygen consumption rates (*J*O_2_) in mitochondria isolated from whole kidney cortex (A & B), and enriched proximal tubule (C & D), distal tubule (E & F), and glomerular (G & H) samples. The conductance of the electron transport chain (ETC) in each of our samples reveals differential ETC conductivity in the enriched samples (I), as well as differing zero-load *J*O_2_, which represents the conductivity of the ETC at minimal bioenergetic resistance (i.e., ΔG’_ATP_ = 0), shows differing bioenergetic profiles of the sieved fractions and the isolated mitochondria, which are differentially impacted by adenine diet (J). *P*: Pyruvate; *G*: Glutamate; *M*: Malate; *D*: ADP; *S*: Succinate; *Omy*: Oligomycin.

To further interrogate these differences, we quantified the ETC conductance (the slope of the *J*O₂/ΔG_ATP_ relationship) and the zero-load *J*O₂ (y-intercept at ΔG_ATP_ ≈ 0). Remarkably, PT mitochondria demonstrated a higher conductance and zero-load *J*O_2_ than both isolated mitochondrial preparations and the other nephron segments under baseline conditions (Figures 4I & 4J)^11^. Adenine feeding impaired both metrics across compartments, with the effect clearest in PT (conductance p = 0.049; zero-load *J*O₂ p = 0.049), while other segments showed non-significant trends (conductance: p = 0.182 mito, p = 0.472 DT, p = 0.425 glom; zero-load *J*O₂: p = 0.112 mito, p = 0.441 DT, p = 0.402 glom). Physiologically, higher PT conductance indicates greater responsiveness of electron flux to changes in energetic demand (steeper *J*O₂ gain per unit decrease in ΔG_ATP_), and the elevated zero-load *J*O₂ reflects a larger “no-load” respiratory capacity. The adenine-induced drop in both measures suggests a diet-induced combined insult, making PT mitochondria both the most capable and the most vulnerable to adenine-induced bioenergetic losses in this model.

### Segmental Regulation of Mitochondrial Membrane Potential Reflects Bioenergetic Demand

Recent studies in tissues like skeletal muscle have demonstrated that ΔΨ_mt_ is not static, but rather modulates in response to shifts in energetic demand^17–19^. These adaptations are often associated with cristae remodeling and heterogeneous cristae membrane potentials to optimize ATP synthesis under differing physiological states^18,19^. To evaluate whether similar mitochondrial membrane potential dynamics are evident in kidney mitochondria, we employed the CK clamp in combination with tetramethylrhodamine methyl ester (TMRM), a potentiometric dye whose fluorescence increases with ΔΨ ^23,24,38,51^. This approach allowed us to assess changes in mitochondrial polarization across a gradient of bioenergetic states, simulating varying workloads by titrating PCr.

Surprisingly, both isolated mitochondria and enriched PT samples exhibited a decrease in TMRM fluorescence as energetic demand decreased, indicated by increasing PCr levels under CK clamp conditions (Figures 5A-D). This decline in ΔΨ_mt_ under low ADP availability suggests a tight coupling between ATP turnover and mitochondrial polarization in PT, where a drop in cellular energy demand leads to a proportional decrease in the proton motive force. This behavior contrasts sharply with oxidative tissues such as skeletal muscle, where ΔΨ_mt_ typically increases during resting conditions to preserve a reserve capacity for rapid ATP production.

**Figure 5.**
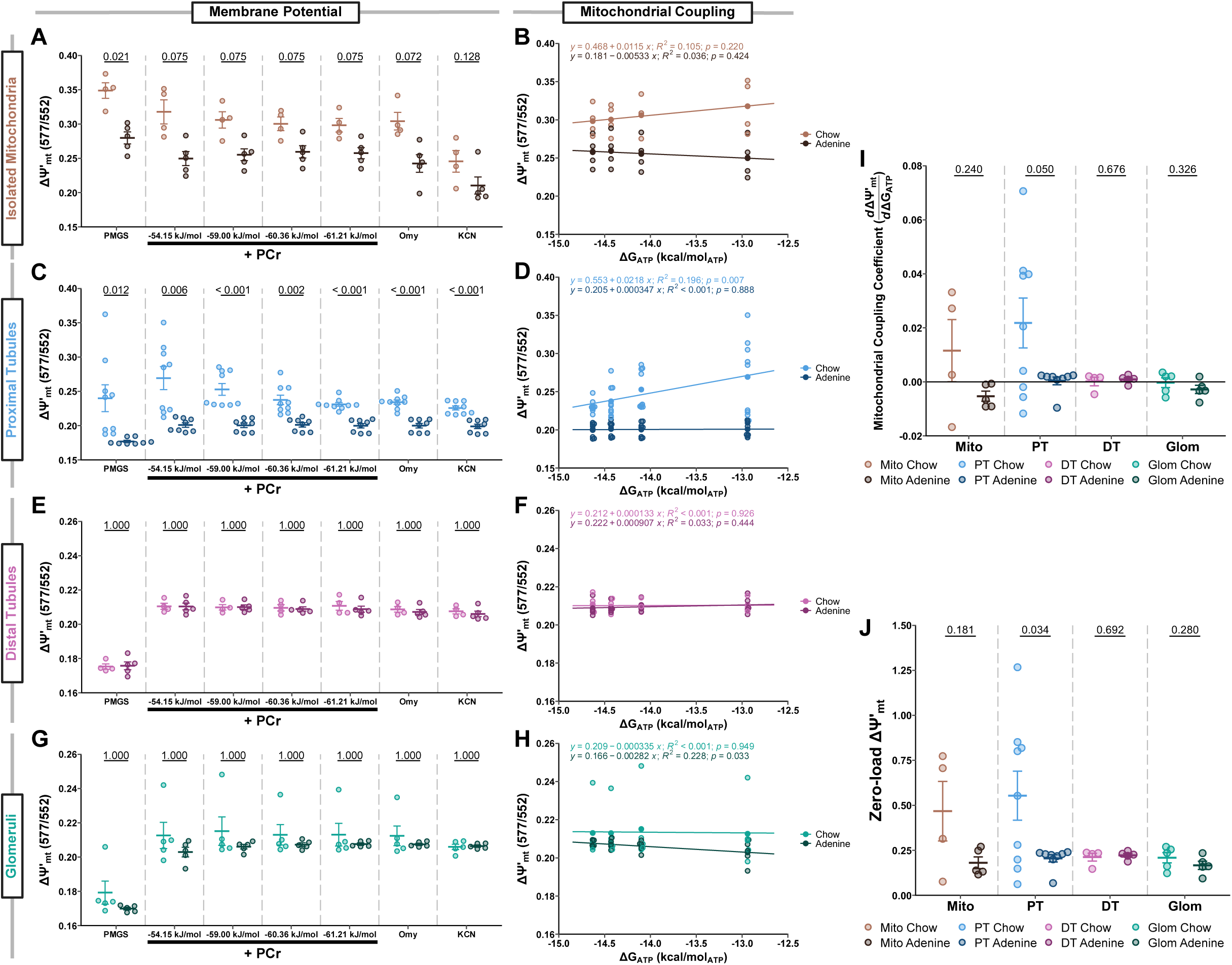
Results of the CK clamp assay on mitochondrial membrane potential (ΔΨ’_mt_) in mitochondria isolated from whole kidney cortex (A & B), and enriched proximal tubule (C & D), distal tubule (E & F), and glomerular (G & H) samples. The coupling coefficient (I), or the slope of the relationship between Gibbs free energy of ATP production and ΔΨ_mt_, outlines the coupling relationship between ΔΨ’_mt_ and ATP production based on the chemiosmotic coupling theory proposed by Peter Mitchell. The zero-load ΔΨ’_mt_, which represents the coupling of ΔΨ’_mt_ to ATP production at minimal bioenergetic resistance (i.e., ΔG’_ATP_ = 0) shows distinctive bioenergetic properties of the isolated mitochondria and the enriched samples, which can also be influenced by adenine feeding (J). *P*: Pyruvate; *G*: Glutamate; *M*: Malate; *D*: ADP; *S*: Succinate; *Omy*: Oligomycin; *KCN*: Potassium cyanide.

This divergent behavior may reflect the distinct physiological roles of these tissues. Skeletal muscle is routinely subject to large, acute swings in ATP demand during transitions from rest to exercise, with energetic demands increasing up to 100-fold during maximal exertion^52,53^. To accommodate this, skeletal muscle mitochondria maintain an elevated ΔΨ_mt_ during rest, likely functioning as an energetic reservoir that can be rapidly mobilized in times of need. This bioenergetic strategy necessitates the preservation of electrochemical potential even during low metabolic activity, ensuring immediate responsiveness to high ATP turnover upon initiation of physical activity.

Conversely, renal PT cells experience fluctuations in energy demand that are less extreme but still dynamic^7,10^, as they are driven by continual solute reabsorption and transport processes that adjust in response to systemic needs. While more modest than those seen in skeletal muscle, these variations still require a finely tuned and adaptable mitochondrial system. Mitochondria in PT may function more like an energetic “current,” dynamically matching ATP production to demand with high efficiency rather than maintaining a stored potential. This tight feedback between energy need tied to continuous solute reabsorption and ΔΨ_mt_ likely optimizes oxidative phosphorylation while minimizing wasteful overproduction of reducing equivalents.

In contrast, DT and Glom samples demonstrated no appreciable changes in TMRM fluorescence across the PCr titration series (Figures 5E-H), suggesting a relatively static ΔΨ_mt_. This observation likely reflects their lower and more stable bioenergetic demands, consistent with their physiological roles in water conservation and filtration barrier maintenance^7,8^. These cell types do not require the same level of rapid energy responsiveness as PT or skeletal muscle, and as such, their mitochondria may favor stability over adaptability. Mitochondria in these segments may operate near maximal capacity even under basal conditions, potentially prioritizing constancy in ATP production and membrane polarization to support prolonged function over acute responsiveness. This static behavior may reflect an evolutionary adaptation to ensure uninterrupted maintenance of essential renal functions, such as osmotic balance and barrier integrity, under a range of metabolic conditions, suggesting a relatively static ΔΨ_mt_.

Although this mode of regulation may be beneficial for maintaining baseline function, it could also predispose these segments to injury under sustained energetic surplus. In conditions such as obesity or high-salt diets, chronic mitochondrial hyperpolarization and reduced capacity for ΔΨ_mt_ adaptation could promote excessive reactive oxygen species (ROS) production, leading to oxidative damage, inflammation, and eventual decline in renal function. These data support a mechanistic framework linking segment-specific mitochondrial behavior to susceptibility in metabolic kidney disease, particularly in distal nephron and glomerular compartments that lack the energetic adaptability observed in PT mitochondria.

To formalize these ΔΨ_mt_ dynamics, we derived two summary metrics from each CK-clamp titration. First, we derived a mitochondrial coupling coefficient (the slope *d*ΔΨ_mt_/*d*ΔG_ATP_), which reports how strongly membrane polarization tracks changes in phosphorylation demand, and second, the zero-load ΔΨmt (the y-intercept at ΔG_ATP_ ≈ 0), which estimates the maximal polarization/reserve when the phosphorylation load is minimal (Fig. 5I–J). In chow controls, PT had the largest coupling coefficient and the highest zero-load ΔΨ_mt_ among compartments, consistent with a system designed to modulate polarization rapidly as transport energetic workload fluctuates. Adenine feeding selectively blunted both the coupling coefficient and zero-load ΔΨ_mt_ (p = 0.050 and p = 0.034, respectively) in PT, whereas isolated mitochondria, DT, and glomeruli showed no significant changes (coupling p = 0.240 mito, p = 0.676 DT, p = 0.326 glom; zero-load p = 0.181 in mito, p = 0.892 in DT,p = 0.280 glom). Physiologically, a higher coupling coefficient reflects greater gain in chemiosmotic potential energy, while a higher zero-load ΔΨ_mt_ reflects a larger polarization reserve available when ATP turnover is clamped low. Thus, the adenine-induced reduction in both metrics indicates that PT mitochondria lose both responsiveness and reserve, aligning with the parallel decreases we observe in ETC conductance and zero-load *J*O₂. Together, these results position PT as the most bioenergetically capable, yet most vulnerable segment to bioenergetic adaptations to external stimuli.

Finally, we plotted *J*O₂ against ΔΨ_mt_ across CK-clamp steps for isolated mitochondria and PT (Fig. S1A–D). The resulting fits were shallow and non-significant, indicating that instantaneous respiration is a weak predictor of membrane polarization in this assay. This is consistent with the indirect coupling between mitochondrial oxygen flux and membrane potential, where membrane polarization is buffered by leak, substrate supply, ion cycling, and ANT/ATP-synthase control. Accordingly, ΔΨ_mt_ should be measured directly and interpreted relative to energetic load rather than inferred from *J*O₂ alone. This does not, however, diminish the value of *J*O₂ as a surrogate of mitochondrial performance, as oxygen consumption remains undeniably central to OXPHOS. Rather, it underscores the need for multidimensional bioenergetic profiling when characterizing mitochondrial function within physiological systems.

### Application to Human Renal Samples

We next extended the sieving–adhesion workflow to human renal cortex obtained from non-implantable deceased-donor kidneys and non-neoplastic (tumor-adjacent) cortex from nephrectomies for clear cell renal cell carcinoma (ccRCC). For human tissue, we adjusted the fractionation to reflect larger nephron dimensions^54–57^: coarse material was removed on a 200 µm mesh; intact glomeruli were collected on a 150 µm mesh (consistent with reported human glomerular size), proximal tubules at 70 µm, and distal tubules at 40 µm (Figure 6 A-D). Pilot testing indicated the mouse digitonin dose was insufficient to permeabilize human preparations; doubling the digitonin concentration restored respiration without excessive cytochrome c sensitivity, and only preparations meeting our outer-membrane integrity criterion were advanced. Using the standard SUIT protocol (due to limited samples from donors), we recorded clear, substrate-responsive oxygen flux from enriched human PT, DT, and glomerular fractions with the expected pharmacologic signatures (Figure 6 D-F). Across Complex I and Complex I+II conditions, ADP-stimulated *J*O₂ ranked PT, DT, and Glom, mirroring the mouse phenotype. Absolute rates showed greater between-sample variability, as anticipated for human tissue (due to possible consequences of donor characteristics and ischemic exposure), but the rank order and inhibitor responses were consistent across sources. These results establish the technical feasibility of segment-resolved mitochondrial bioenergetic assessment in human kidney cortex with minimal front-end adjustments (mesh sizing and increased digitonin), providing a practical bridge from mouse to human and a foundation for application in clinical cohorts.

**Figure 6.**
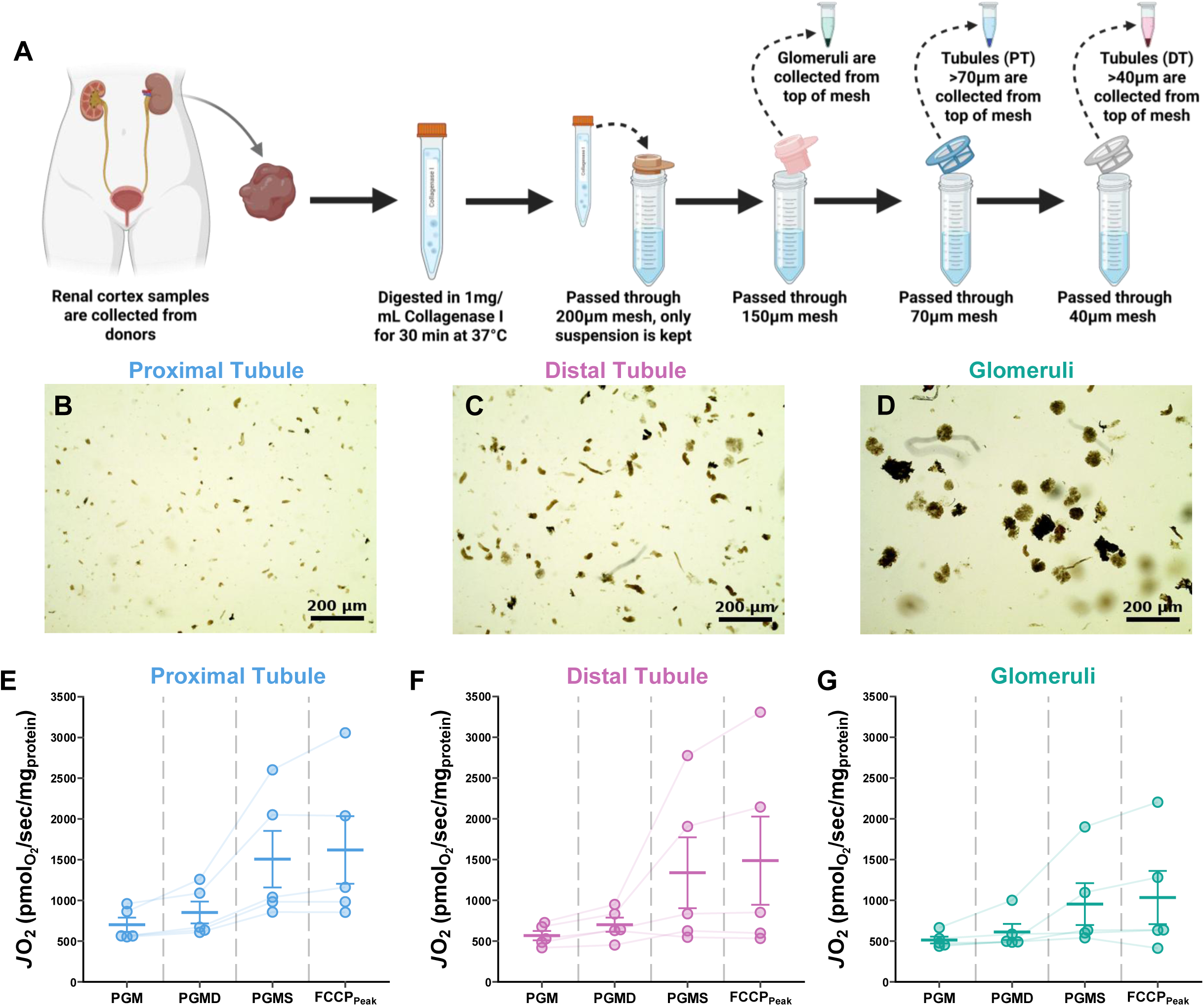
A) Schematic outlining the sieving procedure in kidneys harvested from humans. B) 4X-microscopy images of the enriched tissue samples, indicating the yield and purity of the enriched fractions obtained from this method. C) Initial pilot experiments showing that mitochondrial respiration can be measured in enriched samples from humans. *P*: Pyruvate; *G*: Glutamate; *M*: Malate; *D*: ADP; *S*: Succinate; *FCCP*: Carbonyl cyanide 4-(trifluoromethoxy)phenylhydrazone.

## Discussion

Here, we present a methodological framework that integrates nephron segment enrichment with thermodynamically anchored assessments of mitochondrial function using complementary methods. This approach enables simultaneous evaluation of respiratory flux, electron conductance, and membrane potential within segment-resolved kidney structures. By combining cellular enrichment with control over energetic demand, this method provides a more comprehensive picture of mitochondrial performance under physiologically relevant conditions than is possible with traditional techniques^3,23,42^.

Our results demonstrate that proximal tubule mitochondria exhibit a high degree of energetic adaptability, tightly coupling oxygen consumption and membrane potential to ATP demand across a broad energetic spectrum. This dynamic behavior reflects a capacity to match ATP synthesis to the fluctuating needs of solute reabsorption and active transport. In contrast, mitochondria in distal tubules and glomeruli show markedly limited responsiveness to changes in energetic state, favoring bioenergetic stability over flexibility. These intrinsic differences in energetic plasticity are consistent with the differing physiological roles and energy profiles of each nephron segment and may contribute to segment-specific patterns of vulnerability under pathological conditions.

This distinction in mitochondrial adaptability has significant implications for our understanding of kidney physiology and disease. The high flux sensitivity and tight coupling observed in the proximal tubule suggest a system finely tuned for metabolic efficiency but potentially vulnerable to sudden or sustained stress. Meanwhile, the bioenergetic rigidity of distal segments and glomeruli may confer resilience under normal physiological conditions but may limit adaptive responses during energetic excess or mitochondrial dysfunction. These insights contribute to a growing mechanistic framework that links regional mitochondrial function with renal pathophysiology, including the progression of metabolic kidney disease, diabetic nephropathy, and salt-sensitive hypertension.

More broadly, this integrative platform offers a versatile tool for dissecting cell-type-specific bioenergetic mechanisms in complex tissues. The ability to measure mitochondrial output across a range of controlled ΔG_ATP_ values while maintaining segmental specificity bridges a critical gap between conventional respirometry and organ-level physiological interpretation. In addition to the kidney, this method is readily extensible to other heterogeneous tissues, such as the heart, brain, and liver, where regional mitochondrial differences drive differential responses to stress and disease. Furthermore, this approach is compatible with models of aging, obesity, genetic manipulation, and pharmacologic intervention, making it ideally suited for both discovery research and translational applications.

In summary, the combined approaches of the sieving and adhesion method with high-resolution respirometry and fluorometry, coupled with a SUIT-or CK clamp-based approach introduced here, represent a significant methodological advance for renal mitochondrial physiology. It captures the thermodynamic underpinnings of respiratory control within spatially defined cellular populations, offering both depth and precision in the study of mitochondrial biology. By enabling the detection of subtle, cell-type-specific bioenergetic alterations, this platform lays the groundwork for identifying early signatures of dysfunction and advancing the development of mitochondria-targeted therapies.

## Methods

### Animals

All animal procedures were conducted in accordance with the guidelines established by the National Institutes of Health and approved by the Institutional Animal Care and Use Committee (IACUC) at the University of Utah. Male C57BL/6J mice (8–12 weeks old; Jackson Laboratory, Bar Harbor, ME) were used in all experiments. Mice were housed in a specific pathogen-free facility maintained under standard temperature and relative humidity under a 12-hour light/dark cycle. Animals had ad libitum access to food and water throughout the study unless otherwise specified.

Male mice were specifically selected based on well-documented sex-dependent differences in kidney disease progression, with males exhibiting accelerated onset and more severe manifestations of renal injury in multiple experimental models, including AD-induced nephropathy^32,34^. This design consideration allowed for a more robust and reproducible assessment of mitochondrial dysfunction in response to pathophysiological stimuli.

All animals were group-housed (2–5 mice per cage) in individually ventilated cages with routine health monitoring and environmental enrichment. Mice were randomly assigned to experimental groups and monitored daily for signs of distress or adverse effects. Euthanasia was performed between 8 and 10 AM (to minimize circadian variability) using an intraperitoneal injection of a ketamine (100 mg/kg) and xylazine (10 mg/kg) mixture, followed by thoracic puncture to ensure complete exsanguination and death, in accordance with AVMA guidelines. All efforts were made to minimize animal discomfort and reduce the total number of animals used through careful experimental design and statistical planning.

### Adenine Diet Intervention

For induction of renal injury, mice were fed a high-adenine diet consisting of alternating concentrations of adenine to balance toxicity and survival. Specifically, animals received chow containing 0.15% adenine (w/w) for one week, followed by 0.25% adenine (w/w) for the subsequent week, in a two-week cyclic regimen repeated for the duration of the intervention period. This alternating diet was designed to induce chronic renal stress while minimizing acute toxicity and mortality. Control mice were maintained on standard chow (Teklad Global 18% Protein Rodent Diet, Envigo). All diet formulations were prepared by Envigo and verified for composition.

Body weight, food intake, and signs of discomfort or dehydration were monitored regularly throughout the diet protocol. At the experimental endpoint, mice were euthanized and tissues were harvested for downstream analyses, including histology, nephron segment isolation, and mitochondrial function assays.

### Transmission Electron Microscopy (TEM)

To evaluate mitochondrial ultrastructure, freshly isolated nephron segments were processed for transmission electron microscopy (TEM). Proximal tubule, distal tubule, and glomerular fractions were isolated using the sieving and adhesion method and immediately submerged in fixative solution (1% glutaraldehyde, 2.5% paraformaldehyde, 100 mM cacodylate buffer pH 7.4, 6 mM CaCl2, 4.8% sucrose) and stored at 4°C for 48 hours. Samples were then processed by the Electron Microscopy Core at University of Utah.

Samples underwent 3 × 10-minute washes in 100 mM cacodylate buffer (pH 7.4) prior to secondary fixation (2% osmium tetroxide) for 1 hour at room temperature. Osmium tetroxide as a secondary fixative has the advantage of preserving membrane lipids which are not preserved using aldehyde alone. After secondary fixation, samples were subsequently rinsed for 5 minutes in cacodylate buffer and distilled H2O, followed by prestaining with saturated uranyl acetate for 1 hour at room temperature. After prestaining, each sample was dehydrated with a graded ethanol series (2 × 15 minutes each: 30%, 50%, 70%, 95%; then 3 × 20 minutes each: 100%) and acetone (3 × 15 minutes) and were infiltrated with EPON epoxy resin (5 hours 30%, overnight 70%, 3 × 2-hour 40 minute 100%, 100% fresh for embed). Samples were then polymerized for 48 hours at 60°C. Ultracut was performed using Leica UC 6 ultratome with sections at 70 nm thickness and mounted on 200 mesh copper grids. The grids with the sections were stained for 20 minutes with saturated uranyl acetate and subsequently stained for 10 minutes with lead citrate. Sections were examined using a JEOL 1200EX transmission electron microscope with a Soft Imaging Systems MegaView III CCD camera.

### Histology and Image Analysis

#### Slide Preparation

Kidney samples were fixed in 10% neutral-buffered formaldehyde for 24–48 hours and submitted to the ARUP Research Histology Core Laboratory at the University of Utah. Tissues were embedded in paraffin, sectioned at 4 μm thickness, and stained with Masson’s trichrome to assess morphological changes and fibrotic remodeling.

#### Whole-slide image acquisition

Kidney sections were stained with Masson’s Trichrome (collagen, blue; cytoplasm/muscle, red; nuclei, dark purple). Slides were scanned on a Zeiss AxioScan (ZEISS, Oberkochen, Germany) and converted to pyramidal TIFFs using Fiji. Objective magnification and pixel size were read from image metadata and used to convert pixel counts to physical units (µm and µm²). All downstream analyses operated on 5X-resolution images.

#### Automated Segmentation

In Google Colab (Python 3.10), WSIs were optionally harmonized by blended Reinhard normalization (α = 0.2) and tiled into 1024×1024 px with 256-px overlap. A MONAI 2D U-Net (3 input channels, 7 output classes) was trained on tile/mask pairs to segment glomeruli, tubules, vessels, and background. Slide-level inference used softmax blending with a Hann window; per-pixel labels were assigned by argmax. Class schema, tiling geometry, package versions, and checkpoint paths were logged for reproducibility.

#### Quantification of WSIs

Metrics were computed within semantic masks on the normalized RGB. Collagen burden used Ruifrok–Johnston color deconvolution with mild reconditioning to limit cross-talk; the collagen channel was converted to a concentration map and thresholded per slide by a fixed percentile, followed by speckle removal and a single opening. The glomerular tuft and Bowman’s space were separated from deconvolved intensities along the capsular ridge, and the tuft collagen fraction was reported independently of the periglomerular signal. Tubules were split into instances by a lumen-aware watershed seeded from high-blue (C3) pixels; obvious outliers were filtered by simple shape criteria. For each instance, we report area and standard region-props, lumen area, and pixel-to-µm² conversions from per-slide microns-per-pixel. Outputs consist of per-instance tables (glomeruli, tubules) and lightweight QC overlays; timing and provenance logs accompany each run.

Scripts, training images, and output data are publicly available and accessible on GitHub at https://github.com/stdecker/Trichrome_MONAI_Analysis.

### Nephron Segment Isolation via Sieving and Adhesion

#### Mouse Kidney Specimens and Preparation

Immediately following euthanasia, kidneys were harvested, decapsulated, and placed in ice-cold Hanks’ Balanced Salt Solution (HBSS) without calcium and magnesium. The renal medulla was carefully dissected away to enrich for cortical tissue. Cortices were minced using a sterile single-use razor blade into ∼1 mm³ fragments and incubated in 1 mg/mL collagenase type I (Sigma-Aldrich, SCR103) dissolved in HBSS at 37°C for 20 minutes with gentle agitation. The digested suspension was passed sequentially through nylon mesh strainers (100 μm, 70 μm, and 40 μm) to fractionate nephron segments based on size and adhesion properties.

PT were retained on the 70 μm mesh and rinsed into fresh HBSS. The filtrate was subsequently passed through a 40 μm strainer to collect remaining nephron fragments, which were enriched for DT and Glom. This 40 μm fraction was allowed to settle in a 60 mm tissue culture dish for 2 minutes at room temperature. DT adhered to the plastic surface, whereas Glom remained suspended. The Glom-enriched supernatant was gently removed, filtered again through a clean 40 μm mesh, and washed in HBSS. DT were then carefully scraped from the culture dish and resuspended.

### Human Kidney Specimens and Preparation

Human renal cortex was obtained under institutional approval from two sources: non-implanted deceased-donor kidneys (declined for transplantation) and non-neoplastic (tumor-adjacent) cortex from nephrectomies for clear cell renal cell carcinoma. Tissue was kept on ice, trimmed to remove capsular fat and medulla, minced, and processed using the same sieving– adhesion workflow as for mouse with human-specific mesh cutoffs: coarse material was removed on a 200 µm screen; glomeruli were collected on 150 µm; proximal tubules (PT) on 70 µm; and distal tubules (DT) on 40 µm.

All samples were centrifuged at 1,000 × g for 5 minutes and resuspended in mitochondrial isolation medium (MIM) for subsequent bioenergetic analyses. Aliquots were taken for qPCR-based marker validation and electron microscopy to confirm segmental enrichment and mitochondrial structural preservation.

### Mitochondrial Isolation

Mitochondria were isolated from whole mouse kidney cortical tissue for respirometric and biochemical analyses^28,46^. All tissues were harvested between 9:00 AM and 11:00 AM to minimize circadian variability. Immediately following euthanasia, kidneys were removed, decapsulated, and the renal cortex was rapidly dissected and placed in ice-cold MIM consisting of 300 mM sucrose, 10 mM HEPES, 1 mM EGTA, and 1 mg/mL bovine serum albumin (BSA) (pH 7.4).

Tissue samples were minced with sterile scissors and gently homogenized using a Teflon-glass Potter-Elvehjem homogenizer on ice to preserve mitochondrial integrity. The homogenate was first centrifuged at 800 × g for 10 minutes at 4°C to pellet nuclei and cell debris. The resulting supernatant was collected and centrifuged at 1,300 × g for 10 minutes to further clarify the mitochondrial suspension. This supernatant was then centrifuged at 10,000 × g for 10 minutes at 4°C to pellet intact mitochondria.

The final mitochondrial pellet was resuspended in a minimal volume of fresh MIM buffer depending on the pellet size and experimental application. All mitochondrial preparations were maintained on ice and used immediately for downstream assays, including high-resolution respirometry and ATP production measurements. Protein concentrations were determined using the Pierce BCA Protein Assay Kit (Thermo Fisher Scientific, #23225) following the manufacturer’s instructions. All steps were performed at 4°C or on ice to preserve mitochondrial function and minimize proteolytic degradation.

### Real-time polymerase chain reaction (RT-qPCR)

For gene expression analysis, freshly sieved nephron segments (PT, DT, and Glom) were prepared using the Direct-zol RNA Miniprep Kit (Zymo Research) according to the manufacturer’s protocol, including an on-column DNase digestion step. RNA purity and concentration were assessed spectrophotometrically, and 1 μg of total RNA was reverse transcribed using the iScript cDNA Synthesis Kit (Bio-Rad, Hercules, CA).

Quantitative PCR (qPCR) was performed using SYBR Green-based detection (Thermo Fisher Scientific) in 384-well plates prepared and run by the University of Utah Genomics Core Facility. Expression of kidney injury and fibrosis-related genesPre-validated primer sequences were obtained from the PrimerBank database or prior publications. Relative mRNA levels were normalized to the ribosomal reference gene RPL32 using the ΔΔCt method. Segmental enrichment was validated using primers for segment-specific markers. Nephrin (Nphs1) was used as a glomerular marker (forward: 5’-GACCAGAGGAAGGCATCAAGC-3’; reverse: 5’-GCACAACCTTTATGCAGAACCAG-3’), Slc5a2 (SGLT2) as a proximal tubule marker (forward: 5’-CAGACCTTCGTCATTCTTGCCG-3’; reverse: 5’-GTGCTGGAGATGTTGCCAACAG-3’), and AQP2 as a distal tubule marker (forward: 5’-GCCATCCTCCATGAGATTACCC-3’; reverse: 5’-CGCTCATCAGTGGAGGCAAAGA-3’).

## Mitochondrial Assays

### Substrate-Inhibitor Titration Protocol

#### Isolated Mitochondria

Oxygen consumption by isolated mitochondria was assessed using the Oroboros Oxygraph O2K high-resolution respirometry system (Oroboros Instruments, Innsbruck, Austria), following standard protocols and calibration procedures^28,46^. Mitochondria (75 μg of protein) were resuspended in 2 mL of respiration buffer (Buffer Z) composed of 105 mM MES potassium salt, 30 mM potassium chloride (KCl), 10 mM monopotassium phosphate (KH2PO4), 5 mM magnesium chloride (MgCl2), and 0.5 mg/mL bovine serum albumin (BSA), adjusted to pH 7.2. Buffer Z was equilibrated in the oxygraph chambers at 37°C, while stirring at 750 RPM, prior to sample addition. Following baseline stabilization, mitochondrial respiration was sequentially stimulated by the addition of substrates to support Complex I-and II-linked respiration: 0.5 mM malate, 5 mM pyruvate, 5 mM glutamate, 2 mM ADP, 10 mM succinate, and 1.5 μM FCCP (carbonyl cyanide-p-trifluoromethoxyphenylhydrazone). Oxygen consumption was continuously recorded and analyzed using DatLab software.

#### Enriched Nephron Segments

Mitochondrial oxygen consumption in freshly enriched nephron segments was assessed similarly to isolated mitochondria. Proximal tubules (PT), distal tubules (DT), and glomeruli (Glom) were resuspended in 0.5 mL of Buffer Z (105 mM MES potassium salt, 30 mM KCl, 10 mM KH₂PO₄, 5 mM MgCl₂, and 0.5 mg/mL BSA, pH 7.2) stirring at 750 RPM. Permeabilization was achieved using 5 μL for mouse experiments and 10 μL for human experiments of 0.81 mM digitonin (final concentration: 4 μM and 8 μM), followed by sequential substrate additions to probe electron transport chain function under saturating conditions of hyperoxygenation (200-250 μM O_2_) adapted for tissue diffusion limitations: 5 μL of 0.2 M malate (2 mM), 2.5 μL of 1 M pyruvate (5 mM), 5 μL of 1 M glutamate (10 mM), 5 μL of 0.5 M ADP (5 mM), 5 μL of 1 mM cytochrome c (10 μM), and 10 μL of 0.5 M succinate (10 mM). Maximal uncoupled respiration was induced stepwise with 2 μL of 0.1 mM FCCP (final concentration: 0.4 μM). Approximately 18.75 μg of protein was used per chamber unless otherwise stated. Oxygen consumption was continuously recorded using DatLab software, and only preparations with intact outer mitochondrial membranes (as assessed by the cytochrome c increase of <20% based on pilot data and other published reports^38,58^) were included in analyses.

All measurements were carried out at 37°C to mimic physiological conditions. Care was taken to avoid oxygen limitation, and all chambers were re-oxygenated between runs if needed.

### Mitochondrial ATP Production

#### Isolated Mitochondria

ATP production was quantified fluorometrically using a Horiba Fluoromax 4 fluorometer (Horiba Scientific, USA) by coupling ATP synthesis to NADPH production through an enzymatic cascade as previously described^46^. Assays were performed in using the same substrate conditions as *J*O_2_ at 37°C followed by the addition of 100 μg of protein to each 1 mL assay. The reaction cocktail contained the following components: 2.7 μL of hexokinase (500 U/mL final: 1 U/mL), 3.4 μL of 1 M glucose (final: 2.5 mM), 5.4 μL of 0.05 M AP5A (final: 0.2 mM), 6.8 μL of glucose-6-phosphate dehydrogenase (G6PDH, 500 U/mL final: 2.5 U/mL), and 6.8 μL of 0.5 M NADP⁺ (final: 2.5 mM). Substrates were added to support mitochondrial oxidative phosphorylation: 0.5 mM malate, 5 mM pyruvate, 5 mM glutamate, and 10 mM succinate. Fluorescence was monitored continuously over time.

ATP production was stimulated with sequential additions of ADP to reflect increasing energetic demand, as used previously in our lab (2 μM, 20 μM, and 200 μM). NADPH-linked fluorescence was recorded and used to calculate ATP production rates from the linear portions of the curve.

#### Enriched Nephron Segments

Measurements were conducted using a Horiba Fluoromax 4 fluorometer. Assays were performed in using the same substrate conditions as *J*O_2_ with Buffer Z and digitonin at 37°C, followed by the addition of 100 μg of protein to each 1 mL assay. The reaction cocktail contained the following components: 2.7 μL of hexokinase (500 U/mL final: 1 U/mL), 3.4 μL of 1 M glucose (final: 2.5 mM), 5.4 μL of 0.05 M AP5A (final: 0.2 mM), 6.8 μL of glucose-6-phosphate dehydrogenase (G6PDH, 500 U/mL final: 2.5 U/mL), and 6.8 μL of 0.5 M NADP⁺ (final: 2.5 mM). Substrates were added to support mitochondrial oxidative phosphorylation: 10 μL of 0.2 M malate (final: 2 mM), 5 μL of 1 M pyruvate (final: 5 mM), 10 μL of 1 M glutamate (final: 10 mM), and 20 μL of 0.5 M succinate (final: 10 mM). Fluorescence was monitored continuously over time.

ATP production was stimulated with sequential additions of ADP to reflect increasing energetic demand, following a dosing scheme adapted from skeletal muscle protocols: 50 μL of 0.0005 M (final: 0.025 mM), 50 μL of 0.0005 M (final: 0.05 mM), 10 μL of 0.005 M (final: 0.1 mM), 30 μL of 0.005 M (final: 0.25 mM), and 9.5 μL of 0.5 M (final: 5 mM). NADPH-linked fluorescence was recorded and used to calculate ATP production rates from the linear portions of the curve.

### Creatine Kinase Clamp

#### Isolated Mitochondria

The thermodynamic control of mitochondrial respiration in isolated mitochondria was assessed using the creatine kinase (CK) clamp technique under saturating substrate conditions^47^. Experiments were performed using high-resolution respirometry in Oroboros O2K instruments (Oroboros Instruments, Innsbruck, Austria) at 37°C in 2 mL Buffer Z.

To initiate the CK clamp, 10 μL of 20,000 U/mL creatine kinase (final: 20 U/mL), 2 μL of 0.2 M phosphocreatine (final: 0.2 mM), and 10 μL of 0.5 M ATP (final: 5 mM) were added sequentially. Mitochondria (75 μg protein) were added to the chamber and substrates were introduced to support Complex I and II activity: 5 μL of 1 M pyruvate (final: 5 mM), 5 μL of 1 M glutamate (5 mM), 5 μL of 0.2 M malate (1 mM), and 10 μL of 0.5 M succinate (5 mM).

Phosphocreatine was titrated in a stepwise fashion to establish progressively reduced energetic demand: 6.5 μL of 0.75 M (5 mM), 12 μL (9 mM), and 6 μL (6 mM). To evaluate ATP synthase-dependent respiration, 1 μL of 0.02 mM oligomycin (final: 0.01 μM) was added. Uncoupled respiration was assessed by stepwise addition of FCCP: 1 μL of 0.5 mM (0.5 μM), followed by 1 μL of 1 mM (1 μM), 2 μL (2 μM), and 1 μL (1 μM). Inhibition of respiration was confirmed by the addition of 1 μL of 0.5 mM antimycin A (final: 0.5 μM).

Respiratory flux was recorded continuously using DatLab software and normalized to total protein content.

#### Nephron Segments

Measurements were performed in Oroboros Oxygraph O2K chambers at 37°C using freshly isolated and permeabilized nephron segments (18.75 μg protein in 0.5 mL Buffer Z) under hyperoxygenation (200-250 μM O2). Tissues were permeabilized digitonin followed by sequential reagent additions to establish a CK-clamped energetic environment. The clamp was initiated by addition of 5 μL of 20,000 U/mL creatine kinase (final: 10 U/mL), 2.5 μL of 0.2 M phosphocreatine (final: 1 mM), and 5 μL of 0.5 M ATP (final: 5 mM). To support oxidative phosphorylation, substrates were added as follows: 5 μL of 0.2 M malate (2 mM), 2.5 μL of 1 M pyruvate (5 mM), 5 μL of 1 M glutamate (10 mM), 10 μL of 0.5 M succinate (10 mM), and 5 μL of 1 mM cytochrome c (10 μM).

To impose a progressive energetic gradient and simulate declining ATP demand, phosphocreatine was titrated in successive steps: 3.25 μL of 0.75 M (5 mM), 6 μL of 0.75 M (9 mM), and 4 μL of 0.75 M (6 mM). This approach allowed dynamic assessment of mitochondrial respiratory sensitivity to ΔG_ATP_. After stabilization at the lowest energetic demand, 0.5 μL of 0.02 mM oligomycin (final: 0.02 μM) was introduced to inhibit ATP synthase and quantify proton leak respiration. Finally, maximal uncoupled respiration was determined by addition of 2 μL of 0.1 mM FCCP (final: 0.4 μM).

Oxygen flux was continuously recorded using DatLab software (Oroboros Instruments). Preparations were included only if outer mitochondrial membrane integrity was confirmed by the cytochrome c addition test (<10% stimulation). Respiration rates were normalized to total protein concentration as determined by BCA assay.

##### Mitochondrial Membrane Potential Assay Using TMRM

Mitochondrial membrane potential (ΔΨmt) inwas assessed using a fluorometric assay based on the potentiometric dye tetramethylrhodamine methyl ester (TMRM)^47,51^. All measurements were performed at 37°C in 1 mL quartz or optical glass cuvettes using a Horiba Fluoromax 4 fluorometer (Horiba Scientific, USA) equipped with magnetic stirring. The assay buffer was Buffer Z supplemented with 5 mM creatine monohydrate (freshly prepared). Each well contained 100 μg of protein and was permeabilized with 5 μL of 1 mg/mL digitonin (isolated mitochondria did not have digitonin).

Assay buffer was further supplemented with 0.2 μM TMRM immediately prior to use. A substrate mix was added to support mitochondrial respiration, consisting of 2.5 μL of 1 M pyruvate (5 mM), 2.5 μL of 1 M glutamate (5 mM), 2.5 μL of 0.2 M malate (1 mM), and 10 μL of 0.5 M succinate (10 mM).

The CK clamp was initiated with 5 μL of 20,000 U/mL creatine kinase (final: 10 U/mL), 2.5 μL of 0.2 M phosphocreatine (1 mM), and 5 μL of 0.5 M ATP (5 mM). Phosphocreatine was titrated in a stepwise fashion to progressively increase energy supply: 3.25 μL of 0.75 M (5 mM), 6 μL (9 mM), and 4 μL (6 mM). After establishing steady-state ΔΨmt at each energy state, 1 μL of 0.02 mM oligomycin (final: 0.02 μM) was added to inhibit ATP synthase. Maximal depolarization was induced with 2 μL of 0.5 mM FCCP (0.5 μM) and 1 μL of 0.5 mM antimycin A (0.5 μM).

To determine the appropriate excitation and emission settings, a preliminary excitation scan was performed from 480 to 585 nm, with emission fixed at 600 nm, under four conditions:

(1) buffer only, (2) buffer + mitochondria, (3) buffer + mitochondria + 10 mM succinate, and (4) buffer + mitochondria + 10 mM succinate + 5 μM FCCP. The two excitation wavelengths showing the greatest difference between conditions (3) and (4) were selected for subsequent ratiometric fluorescence measurements. For all assays, optimized excitation/emission settings were 577/552 nm.

Data were expressed as the ratio of TMRM fluorescence under energized versus depolarized states and normalized to protein content.

##### Gibbs Free Energy Calculation

The Gibbs free energy of ATP hydrolysis (ΔG_ATP_) was calculated to quantify the thermodynamic driving force governing mitochondrial energy transduction under CK-clamped conditions^47^. ΔG_ATP_ was derived using the modified Nernst equation:

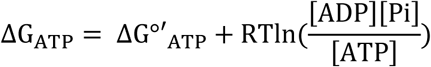

where ΔG°’_ATP_ is the standard free energy change for ATP hydrolysis (typically −30.5 kJ/mol), R is the universal gas constant (8.314 J·mol⁻¹·K⁻¹), T is the absolute temperature in Kelvin (310 K for 37°C), and [ADP], [Pi], and [ATP] are the molar concentrations of ADP, inorganic phosphate, and ATP, respectively.

In the CK clamp system, [ATP] and [ADP] were not directly added in varying concentrations, but instead modulated indirectly via the creatine kinase/phosphocreatine buffer system, according to the equilibrium:

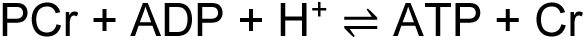

Because the CK reaction is near equilibrium under our assay conditions, the concentrations of ATP and ADP can be inferred from known values of total ATP, PCr, and Cr using the equilibrium constant of the reaction (K′CK ≈ 1.66 × 10⁹ M⁻¹). The assumption of near-equilibrium enables computation of the [ADP]/[ATP] ratio from the known PCr/Cr ratio at each titration point. ΔG_ATP_ values were then calculated across the PCr titration series to generate a force-flux relationship for each biological replicate. Conductance was defined as the slope of the linear regression of *J*O₂ versus ΔG_ATP_.

These values provide a thermodynamic framework for interpreting bioenergetic efficiency and mitochondrial responsiveness across different nephron segments and conditions. ΔG_ATP_ values and associated parameters can be easily calculated using the web-based bioenergetic calculator available at: https://dmpio.github.io/bioenergetic-calculators/^47^.

## Data Analysis and Statistics

Mitochondrial respiratory conductance and membrane potential conductance were calculated by linear regression of oxygen consumption or TMRM signal, respectively, against the corresponding ΔG_ATP_ across phosphocreatine titration steps. The slope of this regression line was used as a measure of respiratory or electrochemical conductance.

Statistical analyses were performed in R version 4.5.1. Unless stated otherwise, tests were two-sided with α=0.05, and data are reported as mean ± SEM. Longitudinal body mass (Diet × Time) was analyzed by two-way ANOVA with Holm-adjusted post hoc contrasts; terminal body, kidney, and liver mass were compared with unpaired two-tailed Student’s t-tests. Histology endpoints were evaluated with linear mixed-effects models (random intercept for individual mice) and estimated marginal means/contrasts via the emmeans package (Holm adjustment). Mitochondrial flux (*J*O₂, *J*_ATP_, CK Clamp) experiments were analyzed using an. Linear trends were fit by ordinary least squares using *lm()* on a per-experiment basis, with mean slopes summarized (e.g., Figures 4 and 5B, D, F, H, I). Model assumptions (normality, homoscedasticity) were evaluated from residuals; when materially violated, variance-stabilizing transformations or the prespecified nonparametric approach were applied. Analyses were conducted using the R packages tidyverse, ggprism, ggbeeswarm, rstatix, ARTool, ggpubr, multcomp, reshape2, ggcompare, mutoss, emmeans, and svglite.

## Acknowledgments

We gratefully acknowledge the support of the University of Utah Health Sciences Core Facilities. Specifically, we thank the Electron Microscopy Core Facility for assistance with ultrastructural analyses, the Cell Imaging Core Facility for support with fluorescence microscopy, ARUP histological services, and the Genomics Core Facility for quantitative PCR services. Their expertise and resources were instrumental in the completion of this study. Schematics in this manuscript were generated using BioRender.

## Author Contributions

S.D. and K.F. conceived the project. S.D., P.C.O., R.H.C., V.L.P., Z.T.S., A.S.K., D.S., L.N., and A.S. performed experiments and collected data. S.D. and L.N. analyzed the data. All authors contributed to the study design and data interpretation. S.D. and K.F. wrote the manuscript. All authors revised the manuscript and approved the final version.

---

**Figure S1.**
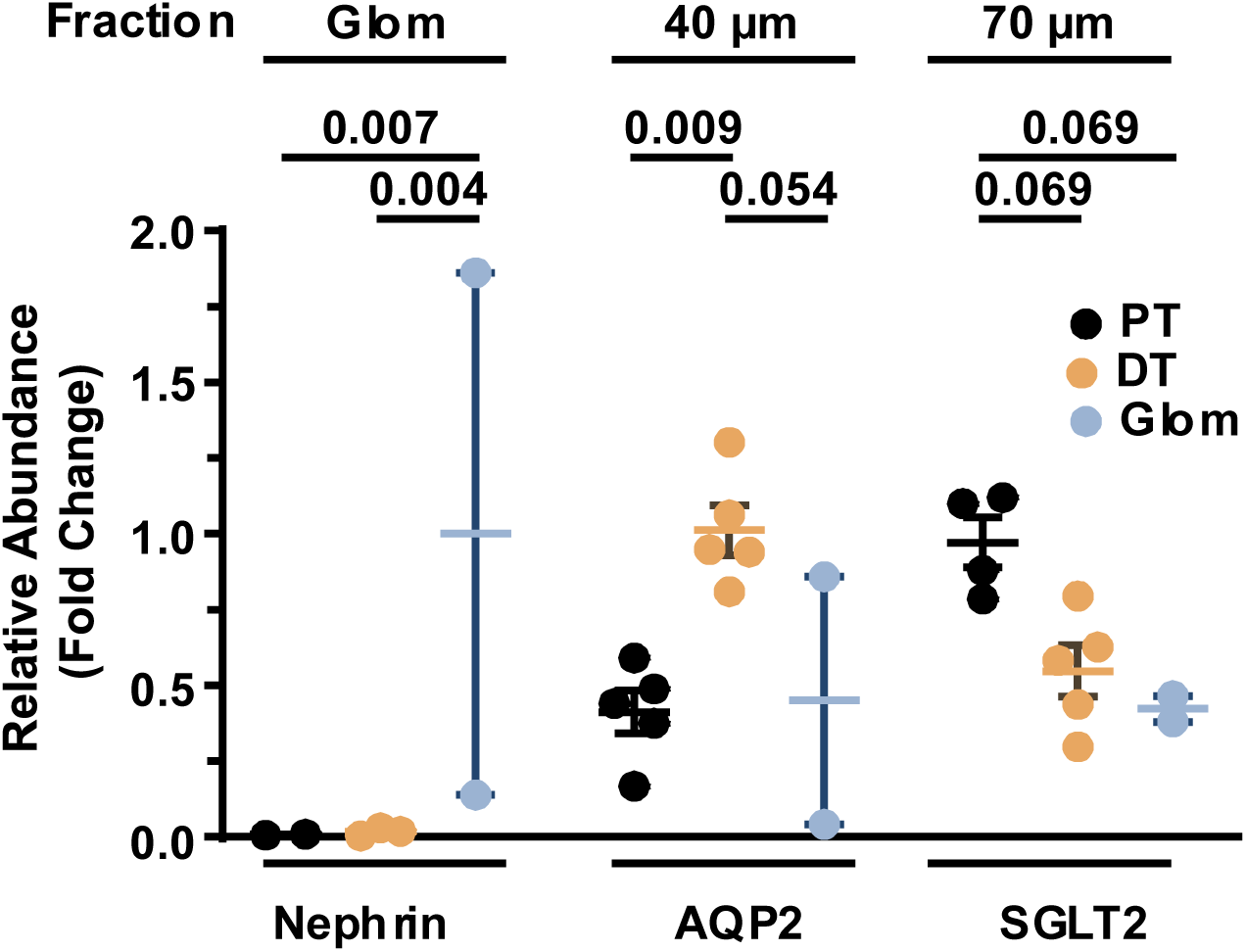
qPCR of our fractions indicating that our samples were yielding enrichment of proximal tubules (PT) at the 70 μm step, distal tubules (DT) at the 40 μm step, and glomeruli in the cell culture dishes. *AQP2*: Aquaporin 2; *SGLT2*: Sodium-glucose cotransporter 2.

**Figure S2.**
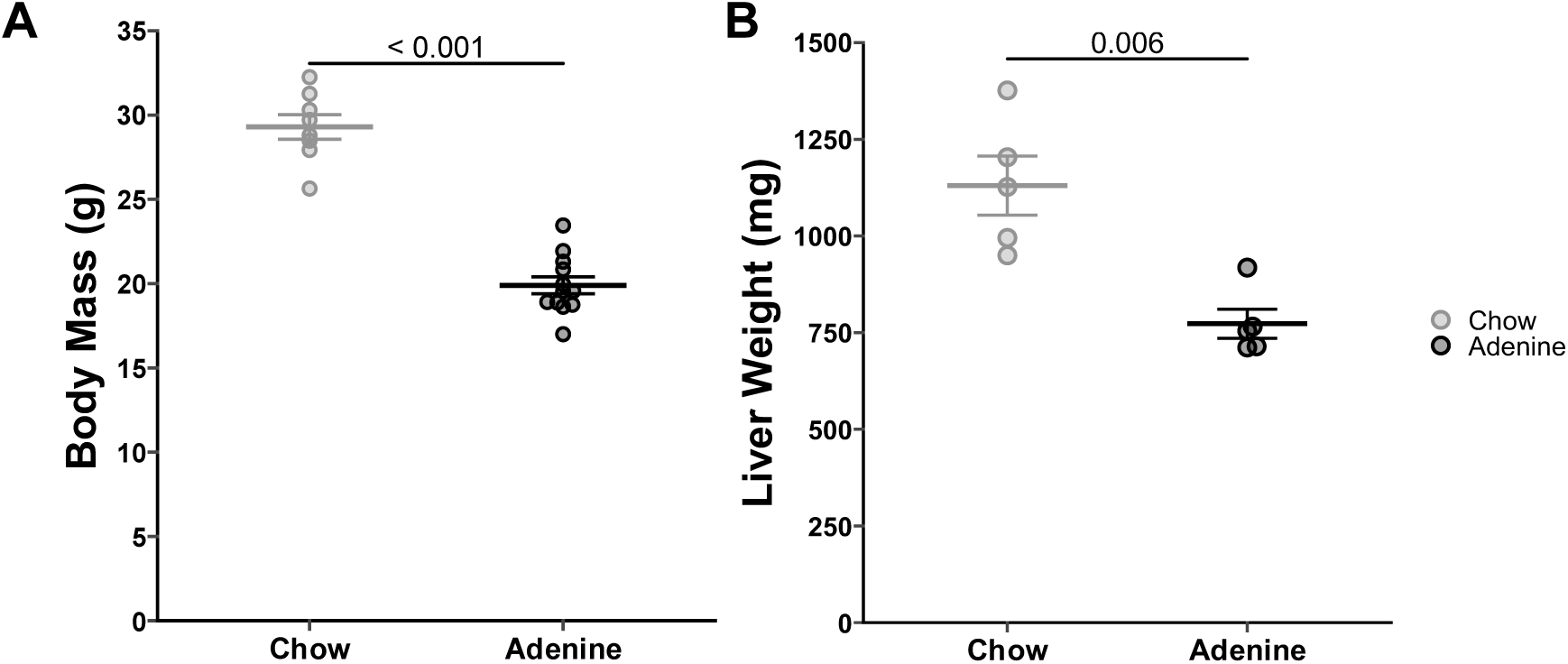
Body mass (A) and liver weights (B) at the time of euthanasia.

**Figure S3.**
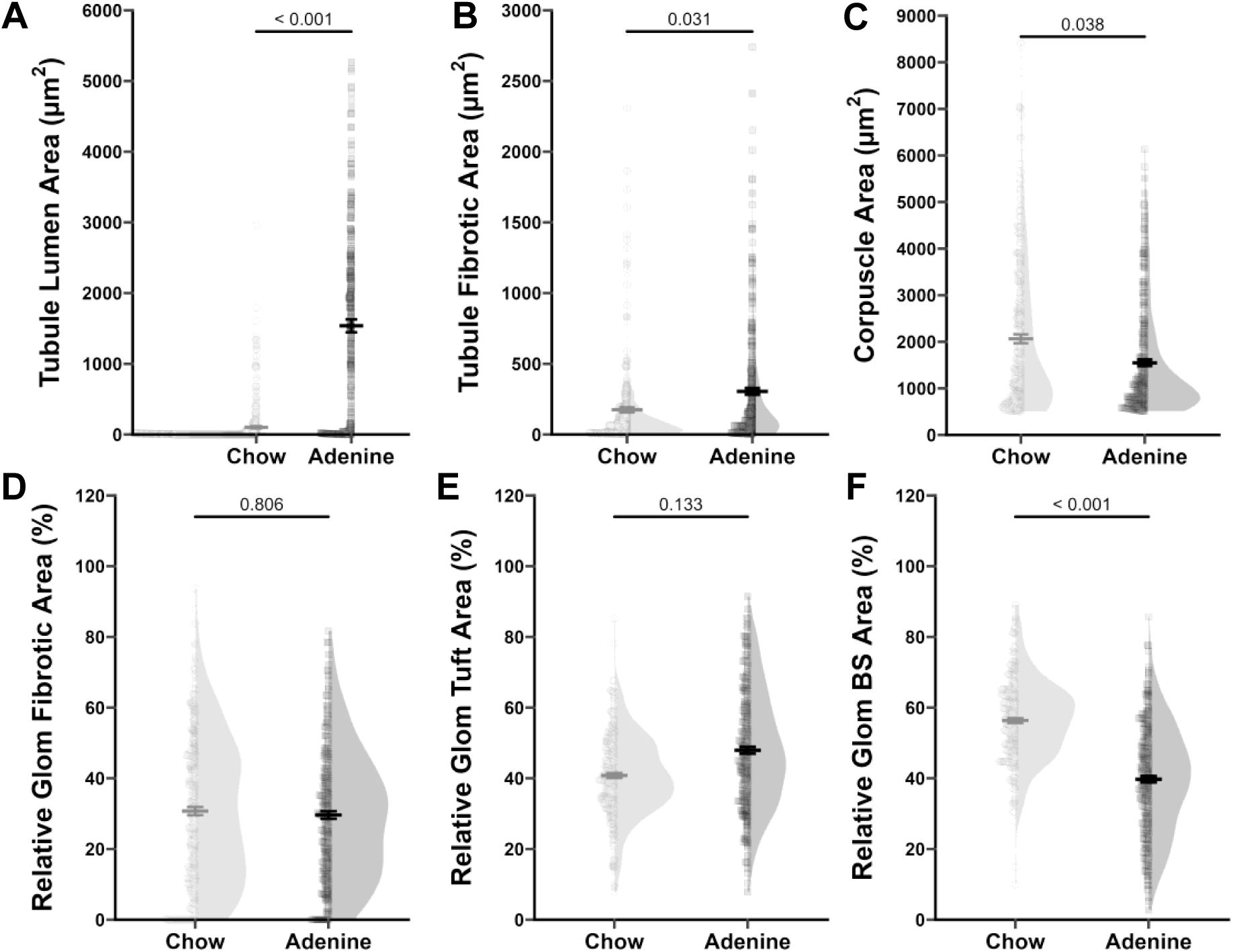
Quantification of hisological parameters obtained from the whole-slide images. Tubule lumen area (A) and fibrotic area (B) were impacted by the adenine diet. Glomerular features, such as corpuscle area and Bowman’s Space (BS) were impacted by adenine diet, but not fibrotic area or tuft area.

**Figure S4.**
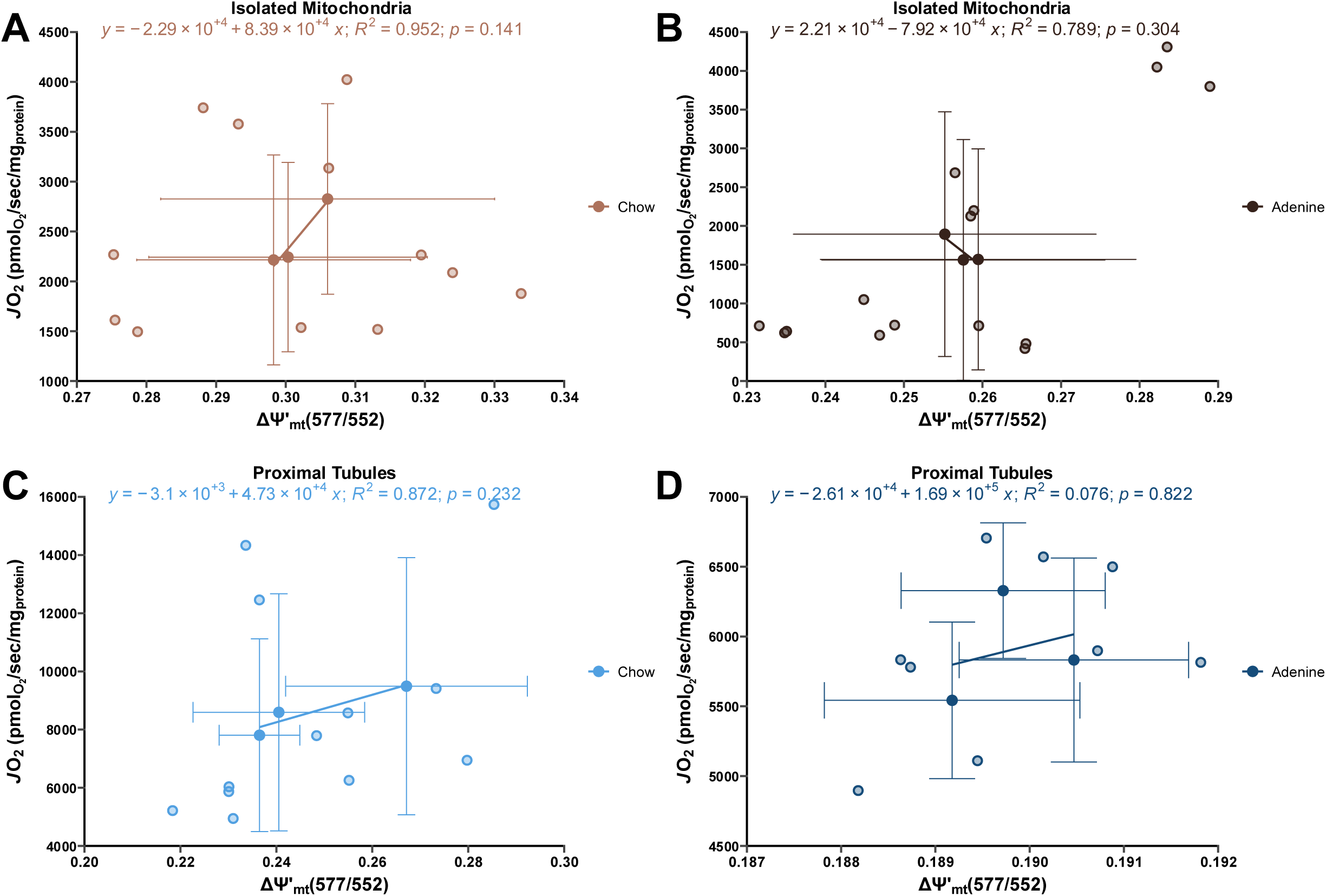
The relationship between oxygen consumption rates (*J*O_2_) and mitochondrial membrane potential (ΔΨ’_mt_) in isolated mitochondria (A & B) and proximal tubules (C & D) shows poor agreement, indicating that *J*O_2_ alone is not a sufficient predictor of ΔΨ’_mt_, likely due to biological uncoupling mechanisms that are known to exist in mitochondrial membranes.

